# Murine norovirus virulence factor 1 (VF1) protein contributes to viral fitness during persistent infection

**DOI:** 10.1101/646281

**Authors:** Constantina Borg, Aminu S. Jahun, Lucy Thorne, Frédéric Sorgeloos, Dalan Bailey, Ian Goodfellow

## Abstract

**Background:** Murine norovirus (MNV) is widely used as a model for studying norovirus biology. While MNV isolates vary in their pathogenesis, infection of immunocompetent mice mostly results in persistent infection. The ability of a virus to establish a persistent infection is dependent on its ability to subvert or avoid the host immune response. Previously, we described the identification and characterization of virulence factor 1 (VF1) in MNV, and established that it acts as an innate immune antagonist. Here, we explore the role of VF1 during persistent MNV infection in an immunocompetent host.

**Methods:** Using reverse genetics, we generated VF1-knockout MNV-3 that contained a single or a triple termination codon inserted in the VF1 open reading frame.

**Results:** VF1-knockout MNV-3 replicated to comparable levels to the wildtype virus in tissue culture. Comparative studies between MNV-3 and an acute MNV-1 strain show that MNV-3 VF1 exerts the same functions as MNV-1 VF1, but with reduced potency. Mice infected with VF1-knockout MNV-3 showed significantly reduced replication kinetics during the acute phase of the infection, but viral loads rapidly reached the levels seen in mice infected with wildtype virus after phenotypic restoration of VF1 expression. Infection with an MNV-3 mutant that had three termination codons inserted into VF1, in which reversion was suppressed, resulted in consistently lower replication throughout a three-month persistent infection in mice, suggesting a role for VF1 in viral fitness *in vivo.*

**Conclusion:** Our results indicate that VF1 expressed by a persistent strain of MNV also functions to antagonise the innate response to infection. We found that VF1 is not essential for viral persistence, but instead contributes to viral fitness in mice. These data fit with the hypothesis that noroviruses utilise multiple mechanisms to avoid and/or control the host response to infection and that VF1 is just one component of this.

## Introduction

With figures upwards of 19 million cases annually, human norovirus (HuNoV) is increasingly becoming the leading causative agent of gastroenteritis [1–4]. Unlike rotaviruses, there are currently no vaccines available against HuNoV, largely due to long-standing difficulties in culturing the virus *in vitro* and the lack of a robust animal model that together have impeded the identification of the correlates of protection. With the identification of murine norovirus (MNV) in 2003 [5], the gap in knowledge on the norovirus lifecycle has been steadily shrinking [6, 7]. MNV has been isolated from wild and laboratory mice, with some research facilities reporting 50-70% seroprevalence [8, 9]. The establishment of reverse genetics systems, the ability to replicate in cultured cells with a tropism for macrophages and dendritic cells, and the availability of a robust homologous small animal model have made MNV a model of choice for studying norovirus pathogenesis [5, 10–12]. MNV-1 was initially discovered as an acute but lethal infection in immunocompromised (STAT1 knockout) mice [5]. Many other strains have since been identified, showing significant differences in their phenotype both *in vitro* and *in vivo* [5, 8, 13, 14]. While some strains cause an acute infection that is cleared within 7 days in immunocompetent mice, other strains are able to persist for up to 9 months [15]. MNV has therefore proven invaluable as a tool to dissect host and viral factors that contribute to viral persistence and the critical role of host factors, such as type III interferons, in the regulation of intestinal infections [7, 16].

Noroviruses are non-enveloped positive-sense single-stranded RNA viruses with a genome ranging from 7.3–7.5 kb [6]. Their small, highly structured genomes include three to four open reading frames (ORFs), with non-structural proteins encoded at the 5’ end and the structural proteins at the 3’ end (Figure 1A). ORF1 encodes a long polyprotein that is post-translationally cleaved by the viral-encoded protease NS6 to produce mature forms of the non-structural proteins, NS1/2–7 [17]. The structural proteins are translated from a subgenomic RNA (sgRNA), with ORF2 encoding the major capsid protein, VP1, and ORF3 encoding the minor capsid protein, VP2. A crucial difference in the genome organisation between HuNoV and MNV is the presence of a fourth and overlapping reading frame within ORF2, found uniquely in MNV [18, 19]. The presence of an alternative reading frame is a common strategy for viruses to maximise the coding capacity of their genome and alternative reading frames often encode accessory proteins that modulate host responses. Indeed, the MNV ORF4 encodes virulence factor 1 (VF1), a mitochondria-localised protein that we have previously shown to be an innate immune antagonist [19]. As yet, whether functional duplication of VF1 exists with another HuNoV protein cannot be excluded, and sequence analysis would indicate that some human sapovirus isolates have an analogous ORF, whose function is yet unknown [20].

**Figure 1.**
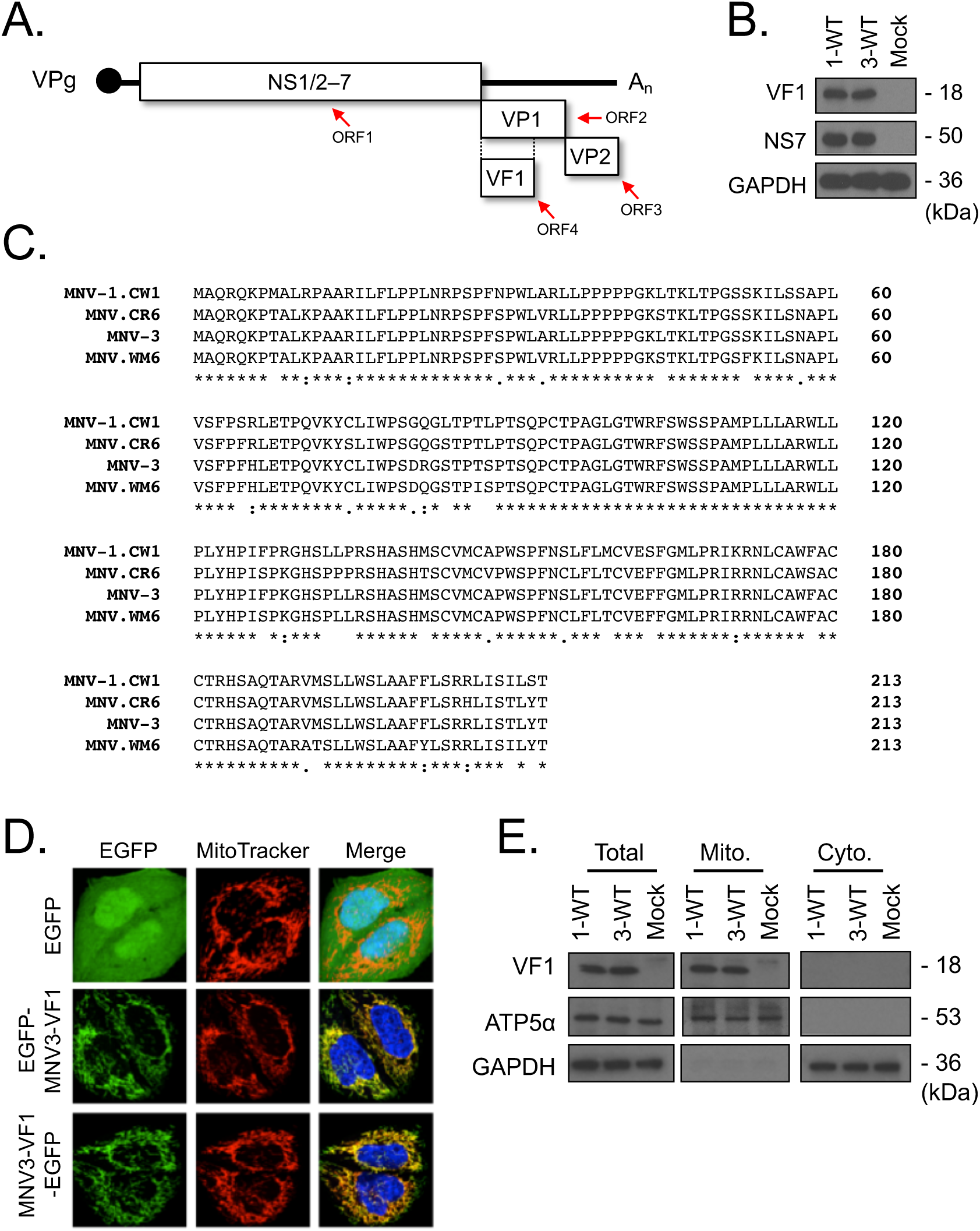
*In vitro* characterisation of MNV-3 VF1. (A) A schematic representation of the MNV genome, showing the four ORFs that encode the non-structural proteins NS1/2–7, VP1, VP2, and VF1. (B) Western blot analysis of VF1 expression following a 12-hour infection with MNV-1 (1-WT) and MNV-3 (3-WT) in RAW264.7 cells (MOI=5). Mock inoculated samples were treated with DMEM. (C) An alignment of VF1 protein sequences from MNV-1, MNV-3, CR6, and WM6. (D) Confocal microscopy of MEFs transfected with plasmids expressing EGFP, or EGFP fused to the N- or C-terminus of MNV-3 VF1. Green indicates EGFP, red indicates MitoTracker, and blue denotes DAPI. (E) Biochemical fractionation of BV-2 cells infected with MNV-1 (1-WT) and MNV-3 (3-WT) (MOI=5). Mock inoculated cells were treated with DMEM. Lysates were collected 12 hours post infection and mitochondrial (mito.) and cytoplasmic (cyto.) fractions were enriched as described in the text.

VF1 is highly conserved amongst MNV strains, including those isolated from wild mice. As it is often the case with viral proteins translated from overlapping reading frames, VF1 bears no significant sequence similarity with any other protein [9, 19]. As an accessory protein, VF1 is not required for viral replication in cell culture but as VF1 expression was systemically restored upon repeated passage of an MNV-1 VF1 knockout virus, its loss likely incurs a fitness cost to the virus. The expression of VF1 was shown to delay expression of key innate immune proteins *in vitro*, most notably interferon-β (IFN-β), revealing its function as an innate immune antagonist [19]. However, the mechanism of action and cellular targets of VF1 have yet to be fully elucidated. *In vivo*, VF1 expression was found to contribute to replication during acute infection in immunocompetent mice and contributed to viral virulence in a lethal model of MNV-1 infection [19].

Host innate immunity is essential in controlling viral replication, placing substantial pressure on viruses to adapt and subvert these defences through a myriad of mechanisms. This dynamic relationship between the host and pathogen determines whether a persistent infection can be established, or if an acute infection is rapidly cleared. Recent studies have shown that early events during infection can dictate the outcome of a persistent infection and the ensuing immune response, as escape of the innate immune response was required to establish a persistent lymphocytic choriomeningitis virus infection *in vivo* [21]. Similarly, the Hepatitis C virus (HCV) protein NS3-4A ablates signalling through RIG-I and MDA5 by cleaving MAVS, permitting HCV to disrupt the innate immune response and establish a persistent infection [22].

In this study, we sought to dissect a potential role for VF1 in viral persistence. To this aim, we examined how the lack of VF1 expression affected the ability of MNV to establish persistent infections in immunocompetent mice. While the introduction of a single stop codon in ORF4 to prevent VF1 expression led to significant reduction in shed virus, expression was restored in infected mice, concomitant with restoration of VF1 expression by the introduction of genetic mutations. Following the introduction of three stop codons in VF1 to suppress this reversion, a consistently reduced level of shedding was observed throughout a three-month persistent infection *in vivo*. Our data confirm that while VF1 is not required for persistence, it nevertheless contributes to viral fitness during persistent MNV infections.

## Methods

### Cell culture and plasmid constructs

The murine microglial BV-2 cell line was kindly provided by Jennifer Pocock (University College London). The cells were maintained in Dulbecco’s modified Eagle’s medium (DMEM, Gibco) supplemented with 10% fetal bovine serum (FBS, Biosera), penicillin (100 units/ml), streptomycin (100 µg/ml), 2 mM L-glutamine and 0.075% sodium bicarbonate (Gibco). Murine macrophage RAW264.7 cells and immortalised murine embryonic fibroblasts (MEFs) were maintained in DMEM supplemented with 10% FBS, penicillin (100 U/mL) and streptomycin (100 U/mL) at 37 °C with 5% CO_2_. Baby hamster kidney cells expressing T7 polymerase (BSR-T7, kindly provided by Karl-Klaus Conzelmann, Ludwig Maximillian University) were maintained in DMEM, supplemented with 10% FCS, penicillin (100 units/mL), streptomycin (100 µg/ml), and 0.5 mg/mL G418. Bone marrow-derived macrophages (BMDMs) were prepared from BALB/c mice as previously described [23].

The previously described full length MNV-3 cDNA clone (pT7:MNV-3) [11], which contains the genome under the control of a T7 promoter, was used for this study. cDNA clones of VF1-mutant viruses that included either a single or a triple stop codon in ORF4 were generated by PCR mutagenesis using KOD hot start DNA polymerase (Novagen). The MNV-3 M1 mutant was generated by inserting a stop codon, through the change of T to A at position 5118. The MNV-3 M3 mutant was generated by inserting three stop codons by the introduction of the mutations T5118A, C5198A, and T5207A. In all cases, the VF1 mutations did not affect the protein coding sequence in the VP1 reading frame.

### Immunoblotting

Unless otherwise stated, BV-2 cells were infected with MNV at a multiplicity of infection (MOI) of 5 TCID_50_/cell, and 12 hours post infection the cells were harvested and lysed in RIPA buffer (50 mM Tris-HCl [pH 8.0], 150 mM NaCl, 1 mM EDTA, 1% Triton X-100, 0.1% SDS). Cell lysates were resolved by SDS-PAGE and then probed using anti-GAPDH (Ambion, AM4300), anti-NS7, anti-ATP5*α* (Abcam, ab14748), or a mouse monoclonal anti-VF1 (Abmart, 4K5) antibodies. The anti-NS7 antibody was described previously [24]. Goat anti-rabbit (Santa Cruz, sc-2030) and goat anti-mouse (Santa Cruz, sc-2031) secondary antibodies conjugated with horseradish peroxidase were used for detection by chemiluminescence using the Amersham ECL Prime Kit (GE Healthcare). Biochemical fractionation of infected cells was performed as previously described [19].

### Reverse genetics

VF1 mutant viruses were rescued using the DNA-based reverse genetics system as previously described [10, 11]. Briefly, BSR-T7 cells were infected with recombinant fowlpox virus expressing T7-polymerase at an MOI of ∼5, and then transfected with 1 μg of the respective cDNA clone using Lipofectamine 2000 (Invitrogen). At 48 hours post transfection, cells were freeze-thawed and clarified lysates were titrated by measuring the 50% tissue culture infectious dose (TCID_50_), measured by looking for signs of cytopathic effect at 4 days post infection. The remaining lysates were used to infect BV-2 cells at an MOI of 0.05 TCID_50_/cell to generate high titre passage 1 or 2 stocks. In all cases, stocks were sequence verified across the VF1 coding region prior to use.

### *In vivo* studies

Three to four-week old male C57BL/6 mice were inoculated with 100 µl of virus containing 1000 TCID_50_ by oral gavage. Mock-infected mice were inoculated with DMEM. Mice were weighed, and faecal and tissue samples were taken at various times post infection. Mock-inoculated mice were euthanised, and tissue samples harvested on day 29 only. Tissue samples included mesenteric lymph nodes (MLN), spleen, duodenum, ileum, caecum, and colon, which were stored in RNA Later (Ambion). This work was carried out in accordance with regulations of The Animals (Scientific Procedures) Act 1986 [25]. All procedures were approved by the University of Cambridge Animal Welfare and Ethical Review Body (AWERB) and the UK Home Office and carried out under the Home Office project licence PPL 70/7689.

### RNA extraction and qRT-PCR

Faecal pellets were accurately weighted, then resuspended and homogenised in PBS at a final concentration of 100 mg of faecal pellet/mL of PBS. Following centrifugation for 5 minutes at 4000 × g, viral RNA (vRNA) was extracted from 100 µl of supernatant using the GenElute mammalian total RNA kit (Sigma-Aldrich), according to the manufacturer’s instructions.

Tissue samples (including MLN, spleen, duodenum, ileum, caecum, and colon) were sliced into 1 mm squares with sterile disposable scalpels. Sterile silica beads and 250 µl lysis buffer were added and tissues were homogenised using the FastPrep-24 homogeniser (MP Biomedicals) at 4 m/s for 1 min. The homogenate was extracted and added to 250 µl of 70% ethanol, and RNA was extracted using the GenElute mammalian total RNA kit (Sigma-Aldrich).

Extracted vRNA from faecal and tissue samples were reverse transcribed using M-MLV RT (Promega) and random hexamer primers (Roche). Viral loads were determined by TaqMan qPCR, as previously described [11]. Briefly, cDNA was added to a mastermix containing 2X Precision MasterMix (Primer Design Ltd) and primers (Sense: 5’-CCGCAGGAACGCTCAGCAG -3’, Anti-sense: 5’-GGCTGAATGGGGACGGCCTG-3’, and TaqMan probe: 5’-ATGAGTGATGGCGCA-3’). Reactions were subjected to 50 cycles consisting of denaturation at 95 °C for 15 seconds, followed by annealing and elongation at 60 °C for 1 minute, using a ViiA7 qPCR machine (AB Applied Biosystems). Viral genome copy number was calculated by interpolation from a standard curve fit generated from known quantities of viral RNA.

### Titration of infectious virus from faecal samples

Viral titres were determined with endpoint dilution assay. Faecal samples collected from inoculated mice were accurately weighed and homogenised in PBS at a final concentration of 100 mg/mL. Resuspended faecal material was then centrifuged at maximum speeds for 5 minutes and 100 µl of supernatant was extracted and subsequently centrifuged at maximum speeds for a further 5 minutes to remove any traces of faecal debris. Purified samples were titrated by TCID_50_ in BV-2 cells as above.

### Caspase activation assay

The activation of caspases 3 and 7 was determined using the Caspase-Glo 3/7 assay kit (Promega), according to the manufacturer’s protocol.

### ELISA

The IFN-β ELISA protocol has been previously described [26]. A monoclonal rat anti-mouse IFN-β antibody (Santa Cruz, sc57201) was used for capture, a polyclonal rabbit anti-mouse IFN-β antibody (R&D Systems, 32401-1) was used for detection, and a goat anti-rabbit-HRP (Cell Signaling Technology, 7074) was used as a secondary antibody. Recombinant mouse IFN-β (R&D Systems, 12400-1) was used as a standard.

### Confocal microscopy

The cells were seeded on cover slips overnight and were transfected with the appropriate plasmids using Lipofectamine 2000 (Invitrogen), according to the manufacturer’s protocol. The cells were incubated for 30 minutes in labelling solution (MitoTracker Red CMXRos, Invitrogen) 24 hours after transfection, washed three times with PBS-T, fixed in 4% paraformaldehyde, washed three times with PBS-T, and then washed twice in PBS. The cover slips were then mounted on slides with Mowiol (Sigma) containing the DAPI nuclear stain. The cells were visualised on the Leica SP5 confocal microscope (Leica Microsystems), and data were analysed with Image J (National Institutes of Health).

### Sequence alignment

Alignment of VF1 protein sequences was carried out using Clustal Omega [27], with sequences from MNV-1.CW1 (accession number: DQ285629.1), MNV-3 (accession number: DQ223042.1), MNV.CR6 (accession number: EU004676.1), and MNV.WM6 (accession number: JN975545.1).

### Statistical analysis

Unless otherwise stated, all statistical analyses were done in Prism v6 (GraphPad), and statistical significance was determined by means of two-way repeated measures ANOVA with Bonferroni multiple comparisons tests. Throughout this manuscript, p>0.05, p≤0.05, p≤0.01, p≤0.001, and p≤0.0001 are indicated by ‘ns’, *, **, ***, and ****, respectively. In all cases, error bars indicate standard error of mean.

## Results

### VF1 is expressed during replication of MNV-3 in tissue culture

The MNV-1.CW1 acute and attenuated strain (MNV-1) originated from the brain of immunocompromised mice and was subsequently plaque purified three times, whereas the persistent MNV-3 strain was isolated from a persistently-infected murine research colony [5, 8]. We have previously demonstrated that VF1, encoded by the MNV-1 ORF4 (Figure 1A), is expressed by 9 hours post infection of murine macrophage cultured cells, that it localises to the mitochondria, and that it antagonises induction of IFN-β [19]. To confirm the expression of VF1 in MNV-3-infected cells, RAW264.7 cells were either mock-infected or infected with MNV-1 or MNV-3 at a high MOI, and harvested at 12 hours post infection. As expected, western blot analysis showed that VF1 expression was readily detected in both MNV-1 and MNV-3 infected cells (Figure 1B).

The mitochondrial localisation of MNV-1 VF1 was previously determined through biochemical fractionation studies, and confocal imaging of GFP-tagged VF1 in transfected cells [19]. Given the high sequence similarity between VF1 proteins from different MNV strains (Figure 1C), we hypothesised that VF1 expressed in MNV-3-infected cells will also localise to mitochondria. To test this, N-terminal or C-terminal MNV-3 VF1 GFP-fusions were constructed and transfected into murine embryonic fibroblasts. As shown in Figure 1D, both the N- and C-terminal MNV-3 VF1 GFP-fusions demonstrated a mitochondrial localisation pattern, akin to that observed for MNV-1 [19]. To examine the localisation of MNV-3 VF1 expressed in the context of viral infection and to rule out any artefacts due to fusion to GFP, we carried out biochemical fractionation on infected murine microglial BV-2 cells. VF1 was detected in the mitochondrial fractions but not in the cytoplasmic fractions in both MNV-1 and MNV-3 infected cells (Figure 1E), indicating a mitochondrial localisation. The host proteins GAPDH and ATP5α were used to confirm enrichment of cytoplasmic and mitochondrial fractions respectively. Taken together, these data indicate that like MNV-1 VF1, MNV-3 VF1 localises to the mitochondria in transfected and infected cells and suggest that its mitochondrial localisation does not require any other viral component.

### VF1 is not required for MNV-3 replication in cell culture

In order to assess a potential role for the VF1 protein in MNV-3 replication, an MNV-3 infectious clone was mutated to introduce one or three stop codons into the VF1 coding frame to produce the mutants M1 or M3 respectively (Figures 2A and 2B). MNV-3 M1 has a single nucleotide substitution (T5118A) that truncates the VF1 protein after 17 amino acids, whereas MNV-3 M3 has three nucleotide substitutions at T5118A, C5198A, and T5207A. VF1 protein is encoded in an alternative reading frame to ORF2, which encodes the major capsid protein VP1, so all mutations were designed to be non-synonymous for VF1 without affecting the overlapping capsid coding sequence (Figure 2B). The yields of infectious MNV-3 M1 and M3 mutants, determined using the previously described DNA-based reverse genetics system [10], were comparable to the WT MNV-3 titres (Figure 2C). To investigate if VF1 was required for MNV-3 replication *in vitro*, replication of MNV-3 M1 and M3 in BV-2 cells was compared to that of WT MNV-3 (Figure 2D). The lack of VF1 had no impact on the ability of MNV-3 to replicate in the immortalised microglial BV-2 cell line, as we have previously seen with MNV-1 [19].

**Figure 2.**
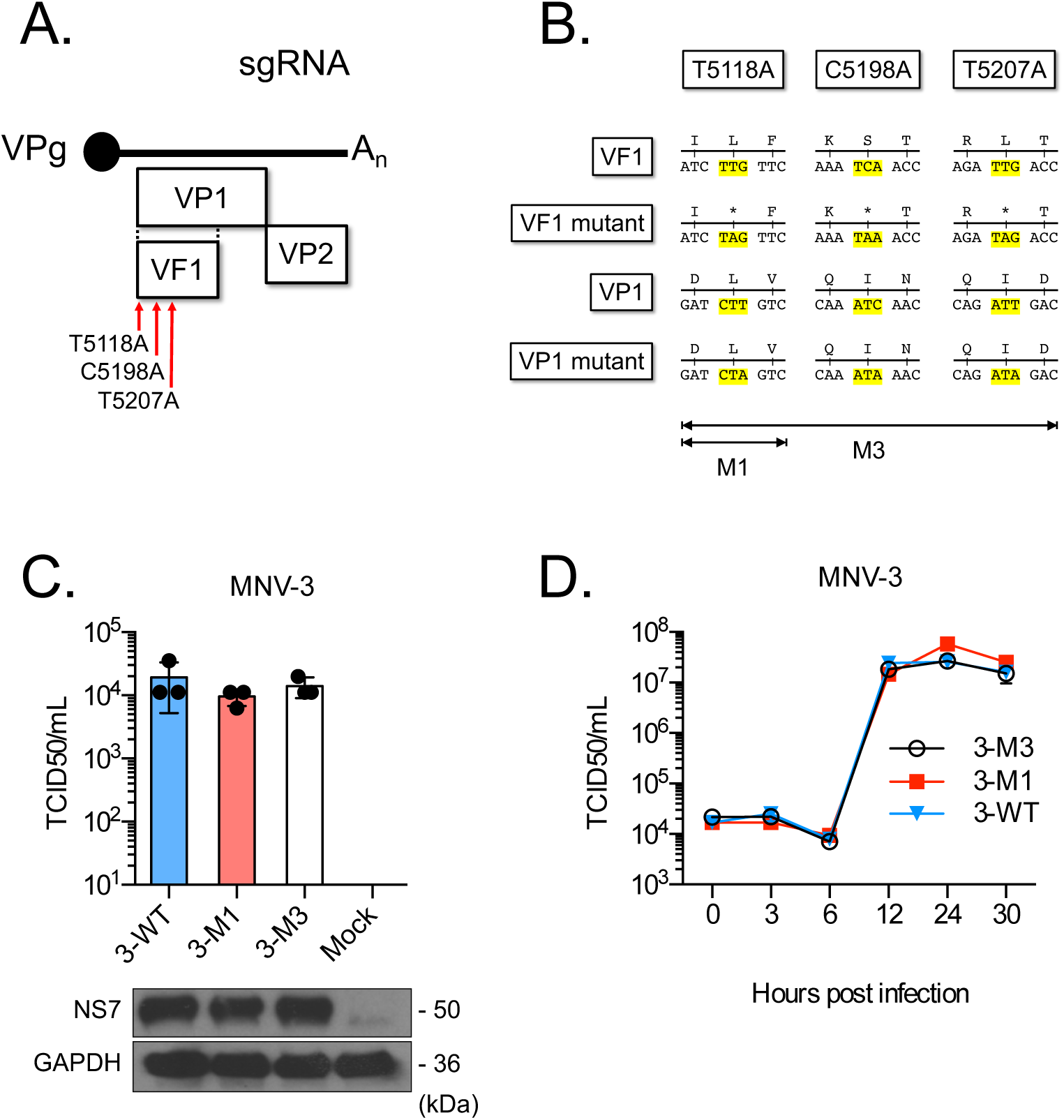
MNV-3 VF1 is dispensable for replication in cell lines. (A) and (B) Schematic representations of the MNV subgenomic RNA indicating sites of stop codon placements for MNV-3 M1 and MNV-3 M3. Stop codon mutations are nonsense for VF1 as they introduce stop codons at position 5118 for MNV-3 M1, and at 5118, 5198, and 5207 for MNV-3 M3. (C) Plasmids encoding full length WT MNV-3 (3-WT), MNV-3 M1 (3-M1) or MNV-3 M3 (3-M3) were transfected into BSRT7 cells. Recovered viruses were titred in BV-2 cells (top panel). Antisera to NS7 and GAPDH were used for western blot analysis (bottom panel). (D) Single-step growth curve analysis. BV-2 cells were infected at an MOI of 5 TCID_50_/cell with WT MNV-3 (3-WT), MNV-3 M1 (3-M1) or MNV-3 M3 (3-M3). Samples were harvested at 0, 3, 6, 12, 24, and 30 hours post infection. Viruses were titred in triplicates in BV-2 cells.

### The MNV-3 VF1 protein inhibits virus-induced apoptosis and interferon induction

We have previously shown that in the absence of VF1, MNV-1-induced apoptosis is increased with enhanced caspase 3/7 cleavage observed in infected cells, indicating that VF1 expression delays apoptosis [19]. However, the induction of apoptosis occurs late in the MNV life cycle, and therefore the impact of the lack of VF1 on virus-induced apoptosis is seen only in the latter stages of the infection [19]. To assess a potential role for VF1 in apoptosis induction during MNV-3 infection, we compared the effect of the lack of VF1 expression on caspase 3/7 activity in MNV-1 and MNV-3 infected cells. We found that the absence of VF1 expression in MNV-3 M1 had only a marginal, but reproducible, impact on the levels of active caspase 3/7 when compared to WT MNV-3 (Figure 3A, top panel). This was in contrast to the more substantial difference observed when comparing WT MNV-1 and MNV-1 M1-infected cells. This impact was not due to any inherent differences in the overall replication rate of MNV-1 and MNV-3 as NS7 expression was comparable (Figure 3A, bottom panel).

**Figure 3.**
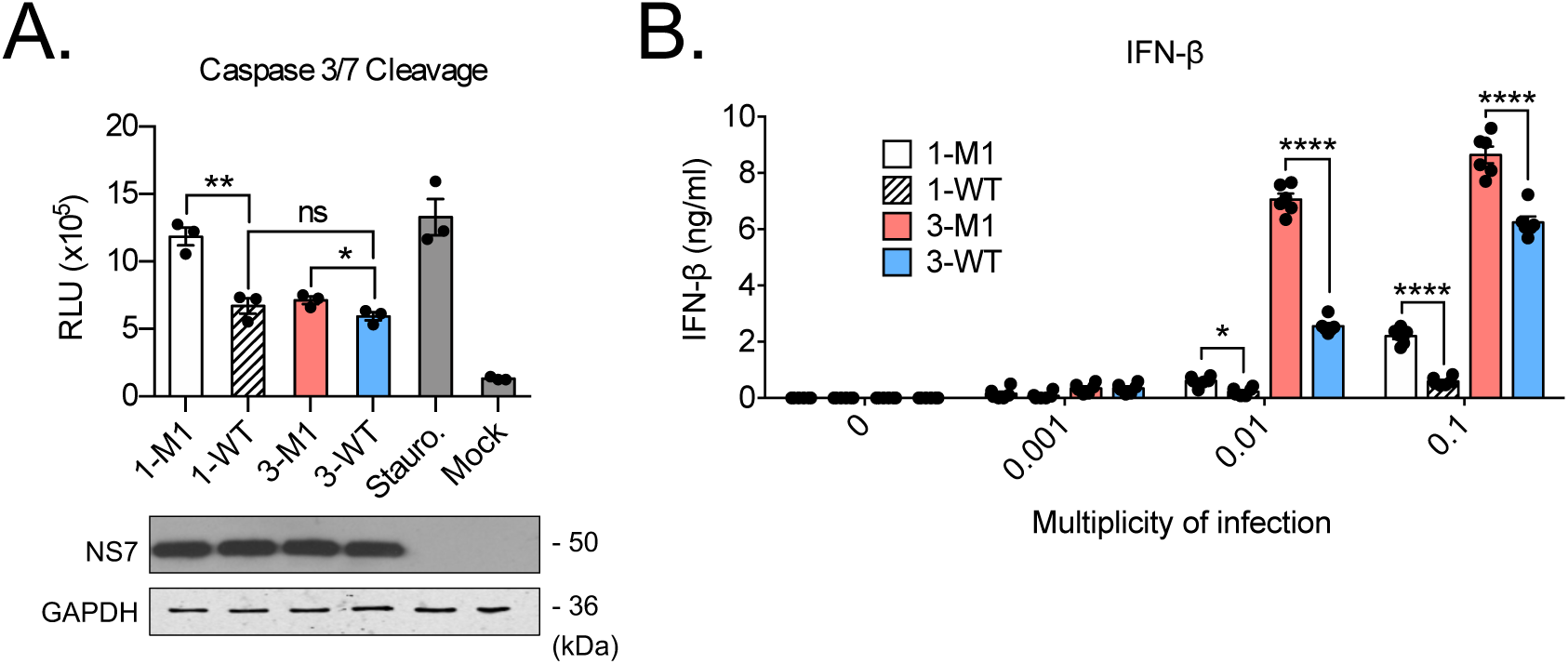
VF1 delays apoptosis and inhibits IFN induction in MNV-3-infected cells. (A) BALB/c BMDMs were infected with WT MNV-1 (1-WT), MNV-1 M1 (1-M1), WT MNV-3 (3-WT) or MNV-3 M1 (3-M1), or treated with staurosporine or DMEM. Caspase 3/7 activity was assayed in triplicates 15 hours post inoculation (top panel). Statistical analysis was carried out using an unpaired t-test. Lysates were analysed by western blotting (bottom panel). RLU, relative luminescence unit. (B) IFN-β ELISA of RAW264.7 cells infected with indicated viruses at different MOI. Cells were harvested at 24 hours post infection. Two-way ANOVA with Bonferroni multiple comparisons tests was used to determine statistical significance. ns=p>0.05; *=p≤0.05; **=p≤0.01; ****=p≤0.0001. ns, not significant.

While VF1 from MNV-1 has been shown to inhibit IFN-*β* induction during infection [19, 28], whether this function is present in MNV-3 is unclear, as reports using VF1 from a variant of MNV-3 that does not persist in immunocompetent mice failed to inhibit RIG-I-dependent induction of a luciferase reporter under the control of the IFN-*β* promoter [28]. Preliminary data from our group however suggest that VF1 can, under certain conditions, inhibit the expression of co-transfected transgenes (not shown), thus complicating the use of reporter-based assays to investigate VF1 function. We therefore explored the role of VF1 in antagonism of IFN-β production during infection using cytokine ELISA. Infection with MNV1 M1 mutant led to significantly increased secretion of IFN-*β* compared to WT MNV1, as expected (Figure 3B). A similar observation was made with MNV-3 confirming that the VF1 protein from MNV-3 also inhibits the innate response. We also noted that the levels IFN-*β* produced from WT MNV-3-infected cells were considerably higher than those infected with WT MNV-1 (Figure 3B).

### Loss of VF1 incurs a fitness cost in MNV-3 replication *in vivo*

Although the MNV-3 VF1 protein was not essential for viral replication in tissue culture, its importance may be more apparent during infection of its natural host. Indeed, some viral accessory proteins that modulate the host response are dispensable for replication in cell culture, but play a role in pathogenicity *in vivo* [29]. To explore this, we orally inoculated immunocompetent mice with sequence-verified WT MNV-3 and MNV-3 M1. As expected based on previous studies [11], no significant effect of WT MNV-3 or MNV-3 M1 infection on weight was observed throughout the duration of the 28-day long experiment (Figure 4A).

**Figure 4.**
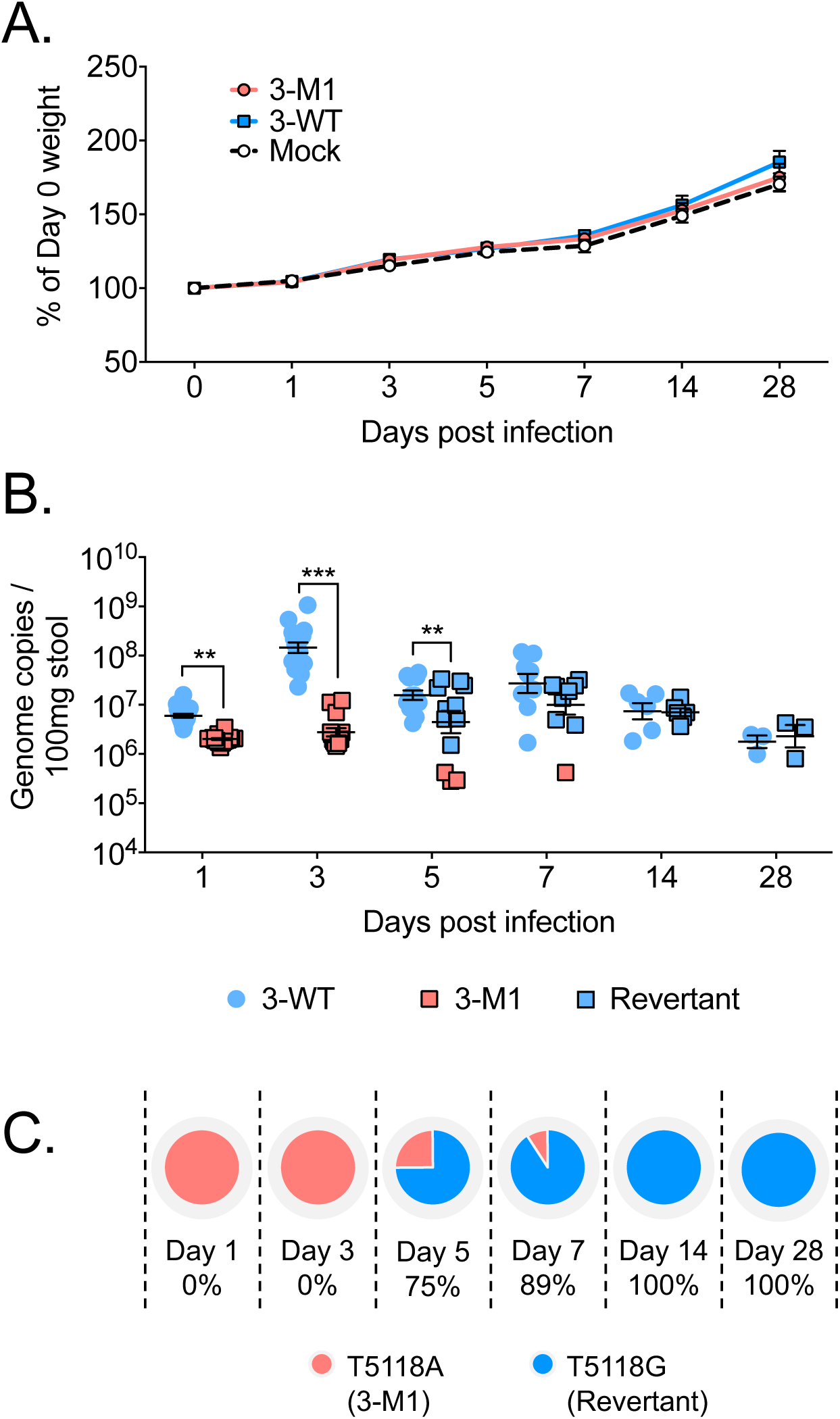
Analysis of MNV-3 VF1 virulence *in vivo*. C57BL/6 mice were orally gavaged with 1000 TCID_50_ of WT MNV-3 (3-WT) or MNV-3 M1 (3-M1). Mock inoculated mice were treated with DMEM. (A) Weights of mice are presented as relative to their day 0 weights (n=8). (B) Viral RNA extracted from faecal samples were quantified using qPCR. Points shaded blue represent VF1 expression (either from WT or revertant viruses). Red points represent absence of VF1 expression. Statistical analyses were calculated using two-way ANOVA with Bonferroni post-tests. (C) Percentages of mice that are shedding revertant viruses after acquiring the A5118G substitution. **=p≤0.01; ***=p≤0.001.

Quantification of viral genome copies from stool revealed that MNV-3 M1 was secreted at significantly lower levels throughout the first 3 days of infection compared to WT MNV-3 (Figure 4B). The greatest difference was observed at day 3 post infection where wildtype titres peaked at 10^6^, but MNV-3 M1 titres were 2-logs lower at 10^4^. These differences began to subside from day 5 onwards, when shedding for MNV-3 decreased and MNV-3 M1 increased. Sequence analysis of the secreted virus at day 5 revealed introduction of a tryptophan at position 17 in VF1, in place of the introduced stop codon. This position is a leucine in WT MNV3, so this mutation represents a phenotypic, non-genotypic reversion of MNV-3 VF1, which occurred in 75% of mice by day 5 (Figures 4B and 4C). In these mice, the reverted virus was shed at comparable levels to WT MNV-3. Conversely, the mice shedding the lowest levels of MNV-3 M1 at day 5 did not have the reverted virus, indicative that absence of MNV-3 VF1 compromises viral fitness and replication in mice, resulting in selective pressure to restore its expression. This effect can also be seen at day 7 whereby 89% of mice shed the revertant virus at comparable levels to WT-infected, while a single mouse was still secreting non-revertant virus at a log lower (Figures 4B and 4C). This phenotypic, but non-genotypic, restoration of MNV-3 VF1 is synonymous in the ORF2 frame, thus not affecting the amino acid sequence of VP1. Taken together, these data indicate that VF1 contributes to viral fitness in MNV-3-infected mice.

To determine the replication kinetics of MNV-3 M1 in tissues, we harvested tissue samples from the MLN, spleen, duodenum, ileum, caecum, and colon of mice infected with WT MNV-3 or MNV-3 M1 for viral genome quantification. As shown in Figures 5A-F, the highest viral copies for both WT MNV-3 and MNV-3 M1 were observed in the MLN at day 7, and in the ileum, caecum, and colon at day 5, while the spleen and duodenum were significantly lower. Similar to the pattern observed for shed virus, MNV-3 M1 was found at significantly lower levels than WT MNV-3 at days 1-3 in MLN, ileum, caecum and colon (Figures 5A, 5D, 5E, and 5F). However, coinciding with the phenotypic reversion that occurs from day 5 that restores VF1 expression, comparable levels of MNV-3 M1 and WT MNV-3 were detected in all tissues with the exception of MLN, where MNV-3 M1 kinetics were still somewhat delayed even after day 5 post infection (Figures 5A-F).

**Figure 5.**
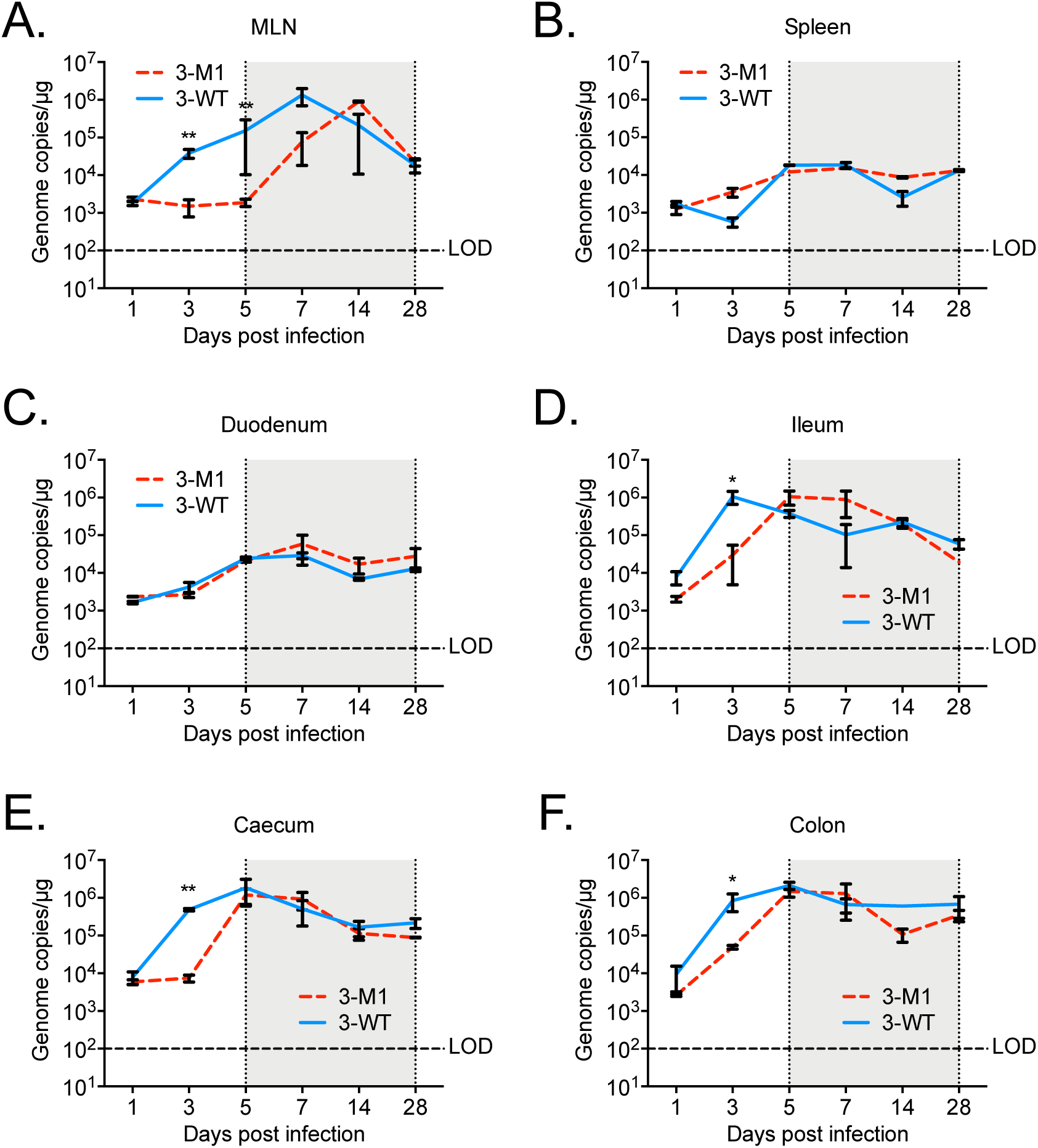
Replication kinetics in tissues. C57BL/6 mice were orally gavaged with 1000 TCID_50_ of WT MNV-3 (3-WT) or MNV-3 M1 (3-M1). Mock inoculated mice were treated with DMEM. Viral RNA was extracted from tissue samples, including (A) MLN, (B) spleen, (C) duodenum, (D) ileum, (E) caecum, and (F) colon, and quantified using qPCR. Statistical analyses were carried out using an unpaired t-test. For WT MNV-3, n=20; for MNV-3 M1, n=20; and for mock, n=5. Shaded area shows the time points from which reversion was seen. *=p≤0.05; **=p≤0.01. LOD, limit of detection.

### MNV-3 VF1 is not required for viral persistence

Advances in the last decade have demonstrated a role for interferon responses in MNV persistence [16]. Treatment with type III IFNs was sufficient for clearance of a persistent strain of MNV [30, 31], and an acute strain of the virus appeared to persist in IFN-*λ* receptor-deficient [32] and conditional type I IFN receptor-deficient mice [33]. Since VF1 is able to antagonise IFN responses, we predicted that it would play a role in viral persistence. The reversion of MNV-3 M1, which had a single stop codon in ORF4, indicates that VF1 likely plays a role in viral fitness, as its absence compromised viral replication in the infected host. However, it also presented a limitation in examining the potential role of VF1 on the establishment of persistence. To prevent the restoration of VF1 expression, we used the triple stop codon mutant MNV-3 M3 characterised in Figures 2A-C. Immunocompetent mice were orally inoculated with sequence verified WT MNV-3 and MNV-3 M3, and stool samples were collected over a 3-month period to quantify viral genome copies with qRT-PCR (Figure 6A). As before, peak shedding was observed at 3 days post infection with WT MNV-3, while MNV-3 M3 is shed at 10-fold lower levels. Shedding of MNV-3 WT and MNV-3 M3 continually decreased over the duration of the study, with MNV-3 M3 consistently shed lower, close to the detection limit. Similar patterns were observed with infectious viral titres from the stool samples, confirming that there was shedding of infectious virus throughout the duration of the study (Figure 6B). Sequence analysis verified the triple stop codons were intact throughout the course of infection, suggesting that VF1 was not necessary for viral persistence but instead contributed to viral fitness during a persistent infection of an immunocompetent host.

**Figure 6.**
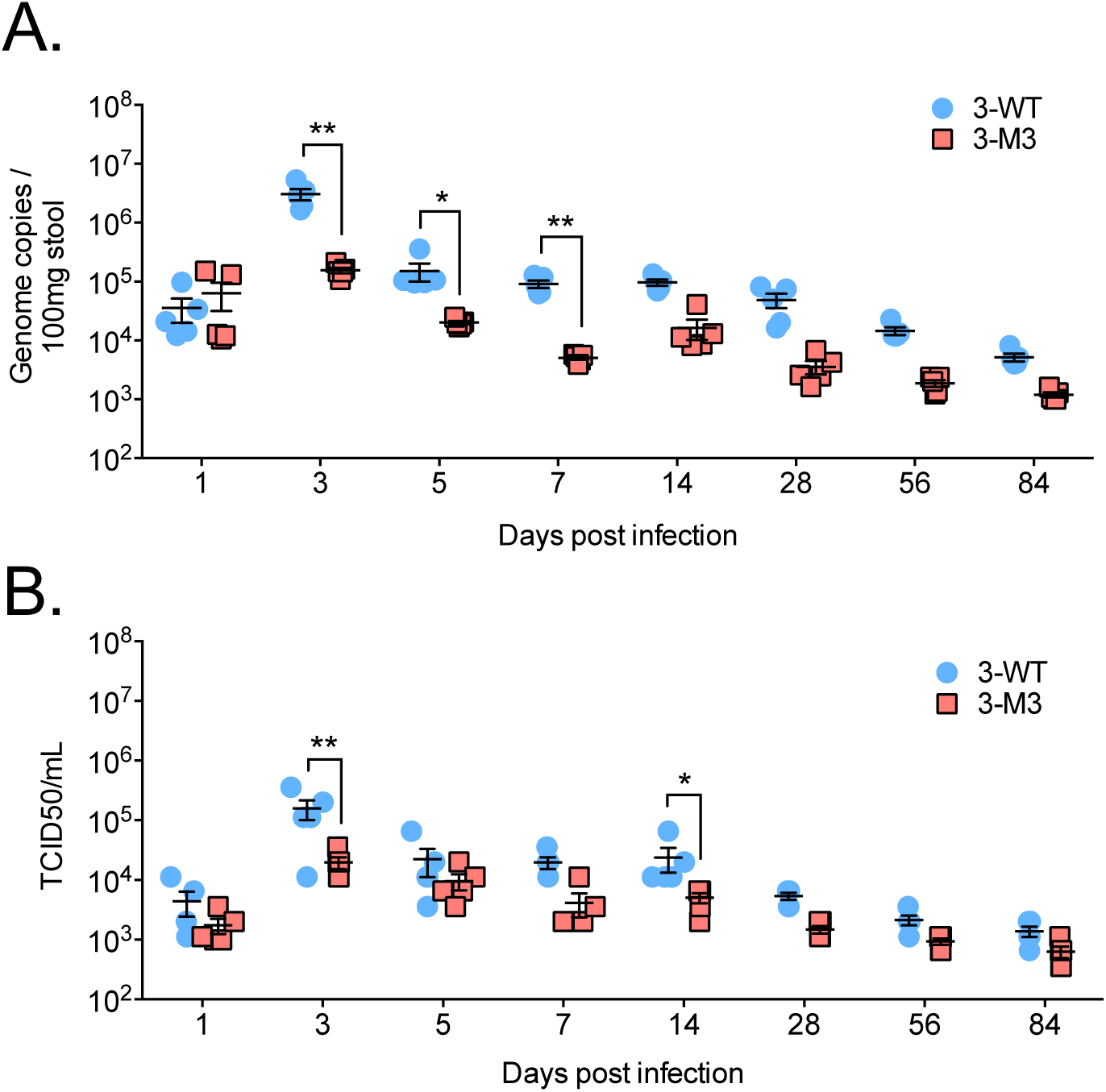
Role of MNV-3 VF1 in persistent infections. C57BL/6 mice were inoculated with WT MNV-3 (3-WT) or MNV-3 M3 (3-M3) at 1000 TCID_50_. Faecal samples were collected over a 3-month (84-day) period. (A) Infectious virus was isolated from faecal samples and titered in RAW264.7 cells. Statistical analysis was carried out using two-way ANOVA with Bonferonni post-test (n=5). (B) Viral RNA from faecal samples was extracted and quantified with qPCR. Statistical analyses were carried out using a two-way ANOVA with Bonferroni post-test (n=5). *=p≤0.05; **=p≤0.01.

## Discussion

The balance between control of infection by host immunity and various mechanisms of subversion by viruses can determine how rapid viral clearance occurs (reviewed in [34, 35]). Since the initial discovery of the acute MNV-1 virus strain in 2003, a number of other MNV strains have been identified that have been shown to persist, including MNV-3 [5, 8, 36]. While MNV-1 is normally cleared in a matter of days, MNV-3 has been shown to persist for up to 9 months [15]. In the context of acute MNV-1 infections, VF1 has been shown to be an innate immune modulator and is important for virulence. In this study we utilised *in vitro* culture systems, reverse genetics and *in vivo* challenge models available for MNV, to compare characteristics and functions of VF1 originating from acute or persistent MNV strains. We showed that the MNV-3 VF1 protein displays the same functions *in vitro* as MNV-1 VF1 and revealed that it is important for viral fitness during persistent infections.

The persistent strain of MNV used in this study, MNV-3, encodes VF1 that is 91% identical to that of the MNV-1 strain at the amino acid level. Sequence variation from the two strains did not affect the mitochondrial localisation of VF1, suggesting that its mitochondrial targeting sequence is conserved, although this has yet to be defined. However, we found differences in the ability of VF1 to delay apoptosis and antagonise innate signaling *in vitro*, with MNV-3 VF1 less potent in both of these activities. Expression levels of VF1 remained similar in assays to compare these functions, so these disparities are likely functional. Although the mechanism of VF1 immune antagonism is yet to be elucidated, the decreased potency of VF1 from MNV-3 may be due to weak binding interactions with an as yet unidentified interacting partner, or inherently decreased functional activity. A comparison of VF1 from multiple MNV strains is therefore warranted and may be informative in identifying key residues important for VF1 function.

Phenotypic differences attributed to VF1 expression were seen in the *in vivo* model of MNV-3 infection that were not observed *in vitro*. Whilst viral replication remains unaffected in tissue culture following knockout of VF1, *in vivo* viral replication is severely compromised during the acute phase of infection, indicating the presence of greater selective pressures imposed on MNV-3 M1 in infected mice. Our data indicate that at 5 days post infection, selection of a revertant virus occurred, most likely due to selective pressures by the host immune system. The same revertant mutation arose in all the mice between 5 and 14 days post infection, reverting from a stop codon to tryptophan, which likely restored VF1 expression. While this mutation was not a genotypic reversion, the revertant virus replicated to comparable levels as the WT virus. We believe this is indicative of a role for VF1 in *in vivo* viral fitness.

To examine a potential role for VF1 in the establishment of persistence, we reduced the likelihood of VF1 restoration by introducing three stop codons in ORF4 to generate the MNV-3 M3 mutant, which replicated at the same levels as WT MNV-3 in tissue culture. Consistent with our findings with the single stop codon mutant MNV-3 M1, there was a significant reduction in replication of MNV-3 M3 in infected mice, compared to the WT virus. VF1 expression remained suppressed throughout the 3-month study as restorative mutations did not occur. At the same time, viral fitness of MNV-3 M3 remained consistently low, as replication levels were maintained comparatively lower than WT MNV-3. Although replicating close to the detection limit, MNV-3 M3 was never cleared from all the mice examined. These findings provide evidence that despite VF1 contributing to viral fitness during persistent infections, it is not required for viral persistence *per se.*

Considerable progress has been made in elucidating both host and viral determinants of persistence in MNV infection. First, comparative studies between the persistent CR6 strain and the acute CW3 strain revealed a genetic determinant of persistence due to a single amino acid change within the 5’ domain of the non-structural protein NS1/2 [37]. Specifically, a change from aspartic acid to glutamic acid at position 94 allows the acute CW3 strain to establish a persistent infection, with the reverse being true for CR6. Second, there is evidence of a critical role played by interferon responses in MNV persistence [16]. Treatment with type III IFNs was sufficient for clearance of a persistent strain of MNV for instance [30, 31], and the acute CW3 strain of the virus persists in IFN-*λ* receptor (IFNLR1)-deficient mice [32], as well as in CD11c-*Ifnar1*-/- mice that have conditional type I IFN receptor-deficient dendritic cells [33]. Lastly, the presence of commensal bacteria in the gut also promotes MNV persistence, as treatment of mice with broad spectrum antibiotics to clear the microbiome also prevents establishment of intestinal persistence of CR6 in a manner dependent on IFNLR1 [38, 39]. There is therefore a complex interplay of viral factors, innate immune defenses (especially type III IFNs), and commensal bacteria that determine persistence, although the exact mechanistic details are still not completely understood. Our data show that inhibition of IFN responses by VF1 does not translate into impairment of persistence but affects viral fitness. The effect of VF1 on type III IFNs induction is yet to be determined, in contrast to NS1/2 for which there is preliminary data that suggest it mediates evasion of type III IFN responses in a strain-specific manner [32]. Nevertheless, the findings described in this paper may open new avenues in investigating norovirus immunopathology through persistence in mice.

## Author contributions

CB, ASJ, LT, FS, DB and IG were all involved in the conceptualization of the study, the interpretation of the results and writing of the manuscript. CB, ASJ, FS and DB were directly involved in the experimental work. IG was responsible for obtaining funding to support the work.

## Acknowledgements

We thank the NIHR Cambridge BRC Cell Phenotyping Hub for providing access to the Leica SP5 confocal microscope. We are also grateful to Jennifer Pocock (University College London) and Karl-Klaus Conzelmann (Ludwig Maximillian University) for generously providing the BV-2 and BSR-T7 cells, respectively, used in this work.

